# Low Perfusion Compartments in Glioblastoma Quantified by Advanced Magnetic Resonance Imaging and Correlated with Patient Survival

**DOI:** 10.1101/180521

**Authors:** Chao Li, Jiun-Lin Yan, Turid Torheim, Mary A. McLean, Natalie R. Boonzaier, Yuan Huang, Jianmin Yuan, Bart RJ Van Dijken, Tomasz Matys, Florian Markowetz, Stephen J. Price

## Abstract

**Background:** Glioblastoma exhibits profound intratumoral heterogeneity in blood perfusion, which may cause inconsistent therapy response. Particularly, low perfusion may create hypoxic microenvironment and induce resistant clones. Thus, developing validated imaging approaches that define low perfusion compartments is crucial for clinical management.

**Methods:** A total of 112 newly-diagnosed supratentorial glioblastoma patients were prospectively recruited for maximal safe resection. Preoperative MRI included anatomical, dynamic susceptibility contrast (DSC), diffusion tensor imaging (DTI) and chemical shift imaging (CSI). The apparent diffusion coefficient (ADC) and relative cerebral blood volume (rCBV) were calculated from DTI and DSC respectively. Using thresholding methods, two low perfusion compartments (ADC_H_-rCBV_L_ and ADC_L_-rCBV_L_) were identified. Volumetric analysis was performed. Lactate and macromolecule/lipid levels were determined from multivoxel spectroscopy. Progression-free survival (PFS) and overall survival (OS) were analysed using Kaplan-Meier and multivariate Cox regression analyses.

**Results:** Two compartments displayed higher lactate and macromolecule/lipid levels than normal controls (each *P* < 0.001), suggesting hypoxic and pro-inflammatory microenvironment. The proportional volume of ADC_L_-rCBV_L_ compartment was associated with a larger infiltration area (*P* < 0.001, rho = 0.42). Lower lactate in this compartment was associated with a less invasive phenotype visualized on DTI. Multivariate Cox regression showed higher lactate level in the ADC_L_-rCBV_L_ compartment was associated with a worse survival (PFS: HR 2.995, *P* = 0.047; OS: HR 4.974, *P* = 0.005).

**Conclusions:** The ADC_L_-rCBV_L_ compartment represent a treatment resistant sub-region associated with glioblastoma invasiveness. This approach was based on clinically available imaging modalities and could thus provide crucial pretreatment information for clinical decision making.

## Introduction

Glioblastoma is a highly aggressive primary tumor in the central nervous system of adults.^1^ Despite advances in treatment, the median overall survival (OS) of patients remains low at 14.6 months.^2^ Inconsistent response is a major challenge in glioblastoma treatment stratification and could be caused by the extensive heterogeneity of this disease. Many genetically distinct cell populations can exist in the same tumor and display diverse treatment response.^3,4^

One of the most fundamental traits of glioblastoma is chronic angiogenesis and elevated perfusion, associated with a more invasive phenotype.^5,6^ However, a potent angiogenesis inhibitor failed to demonstrate consistent benefits in clinical trials of *de novo* glioblastoma.^7^ A possible explanation for this failure is the profound intratumor perfusion heterogeneity in glioblastomas, which is due to aberrant microvasculature and inefficient nutrient delivery. This heterogeneity can give rise to regions within solid tumors where the demand and supply of nutrients is mismatched.^8^ Consequently, the sufficiently perfused sub-regions may hold the advantages for progression and proliferation, whereas the insufficiently perfused sub-regions may have a hypoxic microenvironment.^9^ This microenvironmental pressure may preferentially induce adaptive and resistant clones in the low perfusion sub-regions.^10,11^ There is a rising need to understand the function of low perfusion sub-regions and evaluate their effects on treatment resistance.

Current clinical practice infer the low perfusion regions as the non-enhancing regions within an otherwise contrast enhancing volume on post-contrast T1-weighted structural images, which can lead to non-specific results using the weighted images.^12,13^ Recently, several studies suggested that quantitative imaging features are useful in reflecting intratumor habitats and tumor microenvironment.^14,15^ As such, multiparametric imaging may allow a more comprehensive evaluation of tumor physiology compared to the morphological heterogeneity visualized by structural MRIs.

Here we describe a method for identifying low perfusion compartments in glioblastoma using multiparametric physiological magnetic resonance imaging (MRI). The relative cerebral blood volume (rCBV) calculated from perfusion weighted imaging can measure tumor vascularity^16^. The apparent diffusion coefficient (ADC) estimated from diffusion imaging can potentially provide information about compartments with different cellularity/cell packing^17^. Thus, the two low perfusion compartments we visualized have distinct properties: one compartment with restricted diffusivity that may represent a sub-region adapting to hypoxic acidic conditions^14^, and one compartment with increased diffusivity that may represent a necrotic sub-region with diminishing cellular structure. By combining perfusion and diffusion imaging we studied the metabolic interplay between supply and demand of nutrients in each compartment. Using multivariate survival analysis, we demonstrate that the volume and lactate level of these two compartments are clinically important.

## Methods

### Patient cohort

Patients with a radiological diagnosis of primary supratentorial glioblastoma suitable for maximal safe surgical resection were prospectively recruited from July 2010 to April 2015. All patients had a good performance status (World Health Organization performance status 0-1). Exclusion criteria include history of previous cranial surgery or radiotherapy/chemotherapy, or inability to undergo MRI scanning due to non-MR compatible metallic devices. This study was approved by the local institutional review board. Signed informed consent was obtained from all patients. Patient recruitment and study design was summarized in Figure 1.

**Figure 1.**
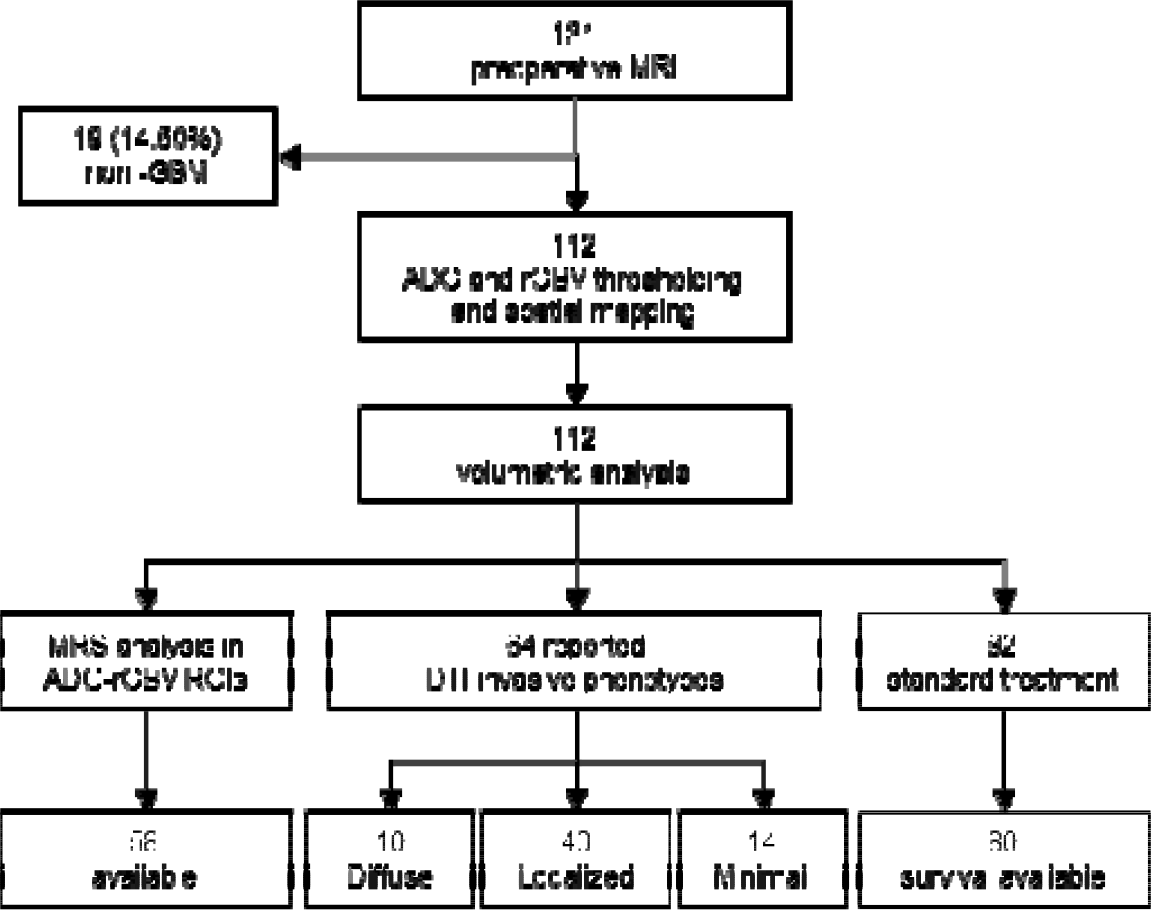
Flow diagram of study design and patient recruitment. Nineteen patients were excluded due to pathological non-glioblastoma diagnosis. Due to the criteria of multiple voxel selection, patients with missing Lac/Cr and ML9/Cr data were excluded in MRS analysis. DTI invasive phenotypes were correlated with the 64 patients overlapping with a previously reported cohort. Patient survival was reviewed retrospectively to exclude psuedoprogression and was only analyzed in those who received standard chemoradiotherapy.

### Treatment and response evaluation

All patients were on stable doses (8mg/day) of dexamethasone. Tumor resection was performed with the guidance of neuronavigation (StealthStation, Medtronic) and 5-aminolevulinic acid fluorescence for maximal safe resection. Extent of resection was assessed according to the postoperative MRI scans within 72 hours as complete resection, partial resection of enhancing tumor or biopsy.^18^

Patients received adjuvant therapy post-operatively according to their performance status. All patients were followed up according to the criteria of response assessment in neuro-oncology (RANO),^19^ incorporating clinical and radiological criteria. Patient survival was analysed for overall and progression-free survival. The latter was made retrospectively in some cases when pseudoprogression was suspected if new contrast enhancement appeared within first 12 weeks after completing chemoradiotherapy.

### MRI acquisition

All MRI sequences were performed at a 3-Tesla MRI system (Magnetron Trio; Siemens Healthcare, Erlangen, Germany) with a standard 12-channel receive-head coil. MRI sequences were acquired as following: post-contrast T1-weighted sequence (TR/TE/TI 2300/2.98/900 ms; flip angle 9°; FOV 256 × 240 mm; 176-208 slices; no slice gap; voxel size 1.0 × 1.0 × 1.0 mm) after intravenous injection of 9 mL gadobutrol (Gadovist,1.0 mmol/mL; Bayer, Leverkusen, Germany); T2-weighted sequence (TR/TE 4840-5470/114 ms; refocusing pulse flip angle 150°; FOV 220 × 165 mm; 23-26 slices; 0.5 mm slice gap; voxel size of 0.7 × 0.7 × 5.0 mm); T2-weighted fluid attenuated inversion recovery (FLAIR) (TR/TE/TI 7840-8420/95/2500 ms; refocusing pulse flip angle 150°; FOV 250 × 200 mm; 27 slices; 1 mm slice gap; voxel size of 0.78125 × 0.78125 × 4.0 mm). Perfusion weighted imaging was acquired with a dynamic susceptibility contrast-enhancement (DSC) sequence (TR/TE 1500/30 ms; flip angle 90°; FOV 192 × 192 mm; FOV 192 × 192 mm; 19 slices; slice gap 1.5 mm; voxel size of 2.0 × 2.0 × 5.0 mm;) with 9 mL gadobutrol (Gadovist 1.0 mmol/mL) followed by a 20 mL saline flush administered via a power injector at 5 mL/s. DTI was acquired before contrast enhanced imaging, with a single-shot echo-planar sequence (TR/TE 8300/98 ms; flip angle 90°; FOV 192 × 192 mm; 63 slices; no slice gap; voxel size 2.0 × 2.0 × 2.0 mm). Inline ADC calculation was performed during DTI acquisition from using b values of 0–1000 sec/mm^2^. Multivoxel 2D ^1^H-MRS chemical shift imaging utilized a semi-LASER sequence (TR/TE 2000/30-35 ms; flip angle 90°; FOV 160 × 160 mm; voxel size 10 × 10 × 15-20 mm). PRESS excitation was selected to encompass a grid of 8 rows × 8 columns on T2-weighted images.

### Image processing

For each subject, all MRI images were co-registered to T2-weighted images with an affine transformation, using the linear image registration tool (FLIRT) functions ^20^ in Oxford Centre for Functional MRI of the Brain (FMRIB) Software Library (FSL) v5.0.0 (Oxford, UK).^21^

DSC data were processed and leakage correction was performed using NordicICE (NordicNeuroLab, Bergen, Norway). The arterial input function was automatically defined and rCBV was calculated in NordicICE. DTI images were processed with a diffusion toolbox (FDT) in FSL, ^22^ during which normalization and eddy current correction were performed. The isotropic component (p) and anisotropic component (q) were calculated using the equation as previously described.^23^

MRS data were processed using LCModel (Provencher, Oakville, Ontario) and the concentrations of lactate (Lac) and macromolecule and lipid levels at 0.9 ppm (ML9) were calculated as a ratio to creatine (Cr). All relevant spectra from MRS voxels of interest were assessed for artefacts using previously published criteria.^24^ The values of the Cramer–Rao lower bounds were used to evaluate the quality and reliability of MRS data and values with standard deviation (SD) > 20% were discarded.^24^

### Regions of interest and volumetric analysis

Tumor Regions of interest (ROIs) were manually segmented using an open-source software 3D slicer v4.6.2 (https://www.slicer.org/) ^25^ by a neurosurgeon with > 7 years of experience (××) and a researcher with > 4 years of brain tumor image analysis experience (××) and reviewed by a neuroradiologist with > 8 years of experience (××) on the post-contrast T1WI and FLAIR. The interrater variability of the ROIs was tested using Dice similarity coefficient scores. For each individual patient, ROIs of normal-appearing white matter (NAWM) were manually segmented from the contralateral white matter and used as normal control. The region is far from the lesion and has no perceivable abnormalities.^26^ All images were normalized by the mean value in NAWM.

ADC-rCBV ROIs were further generated using quartile values in Matlab (v2016a, The MathWorks, Inc., Natick MA). The procedure for identifying these two compartments is illustrated in Figure 2. Firstly, ADC and rCBV values were obtained from each voxel within the contrast-enhancing (CE) ROI delineated on the post-contrast T1-weighted images and pooled together as described previously.^26^ The lowest quartile of the pooled rCBV values (rCBV_L_) were interpreted as low perfusion regions. Then the first quartile (ADC_L_) and last quartile (ADC_H_) of ADC map were respectively overlaid on rCBV_L_ maps. Finally, two intersections of ADC_L_-rCBV_L_ and ADC_H_-rCBV_L_ ROIs were obtained. Other regions within CE outside the two ADC-rCBV ROIs were taken as abnormal controls (CE control, CEC). Raw volumes of ROIs were calculated using the function of fslmaths in FSL.^21^ Proportional volumes (%) of two ADC-rCBV ROIs were calculated as the ratio of the raw volumes to CE volume.

**Figure 2.**
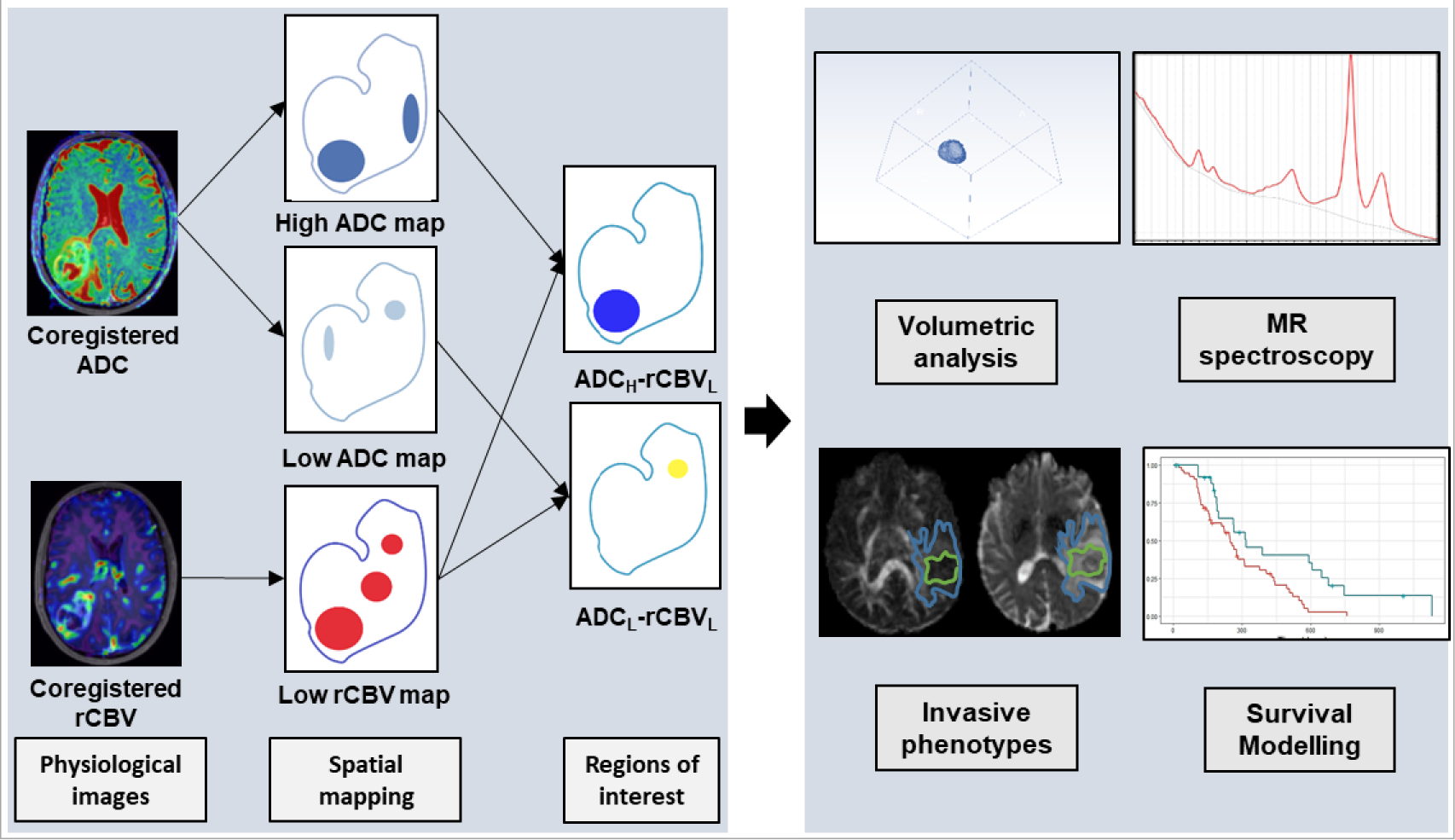
Illustration of the pipeline to identify two ADC-rCBV compartments. Both ADC and rCBV maps are coregistered to the T2 weighted images and tumor regions are segmented manually. Low perfusion tumor regions are partitioned using a quartile threshold. Similarly, two ADC sub-regions are partitioned using high and low ADC quartile thresholds respectively. The spatial overlap between the thresholded rCBV and ADC maps defined two compartments ADC_H_-rCBV_L_ and ADC_L_-rCBV_L_. MR volumetric and metabolic analyses of both compartments are performed and interrogated in invasive tumor phenotype and patient survival analysis models.

### Multivoxel MRS processing

Due to the difference in spatial resolution between MRS and MRI, voxels from T2-weighted MRIs were projected onto MRS according to their coordinates using MATLAB. Thus, the metabolic status in each voxel could be matched to the voxels in the MRI derived physiological maps, which were coregistered to T2-weighted images. The proportion of T2-space tumor voxels occupying each MRS voxel was calculated, and only those MRS voxels that were completely within the delineated tumor were included in further analyses. The weight of each MRS voxel was taken as the proportion of the ADC-rCBV compartments in that MRS voxel. The sum weighted value was used as final metabolic value of the tumor ROI. This method provides an objective method for MRS voxel selection. The selection criteria for MRS data are demonstrated by Figure 3.

**Figure 3.**
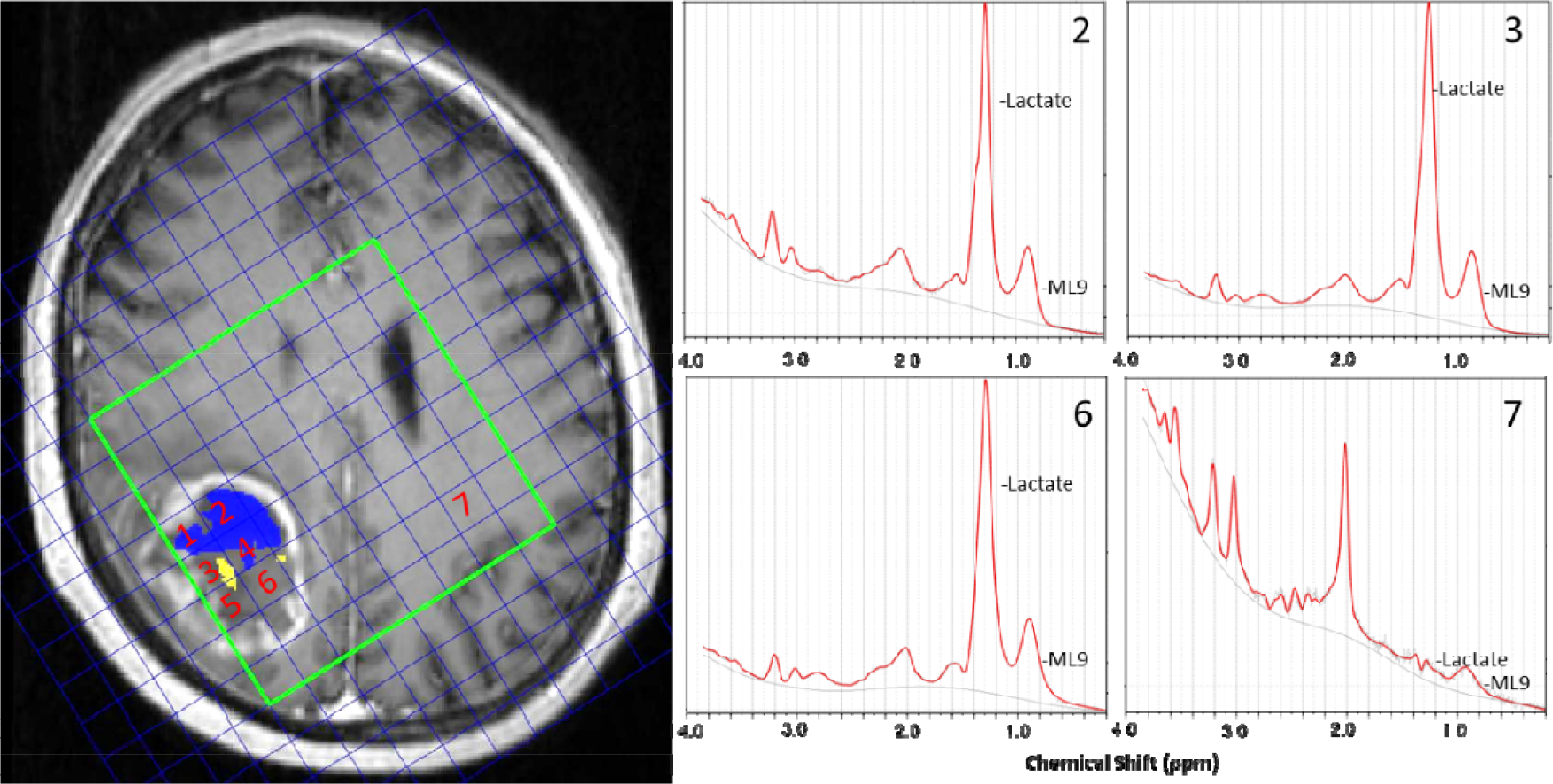
Illustration of multiple voxel MRS analysis. Left: the selection criteria. The T2-space pixels are projected to MRS space according to their coordinates. The proportion of T2-space tumor pixels occupying each MRS voxel is calculated. A criterion is applied that only those MRS voxels are included when this voxel is completely within delineated tumor. In this case, grid 1-6 met the criteria. A weighted average metabolite content for each region is calculated from the metabolite content of each voxel within it weighted by that voxel’s percentage of (ADC_H_-rCBV_L_ [blue]: grid 1,2 and 4 are counted; ADC_L_-rCBV_L_ [yellow], grid 3 and 5 are counted; abnormal control [CEC]: grid 1-6 are counted). Right: Example spectra of ROIs. Each spectrum corresponds to the grids on the left. Grid 2: lactate/Cr ratio 13.2, ML9/Cr ratio: 10.4; grid 3: lactate/Cr ratio 28.4, ML9/Cr ratio: 22.7; grid 6: lactate/Cr ratio 8.9, ML9/Cr ratio: 16.6; grid 7 (NAWM): lactate/Cr ratio 0.37, ML9:1.36.

### DTI invasive phenotypes

To further explore the relation between hypoxic compartments and tumor invasiveness, we investigated DTI invasive phenotypes of 64 patients which overlap with a previously reported cohort and have been correlated to isocitrate dehydrogenase (IDH) mutation status.^27^ Three invasive phenotypes were classified using previously described criteria^28^ based on the decomposition of diffusion tensor into isotropic (p) and anisotropic components (q): (a) diffuse invasive phenotype; (b) localized invasive phenotype; and (c) minimal invasive phenotype. (Figure S1).

### Statistical analysis

All analyses were performed with RStudio v3.2.3. Continuous variables were tested with Welch Two Sample t-test. MRS data or tumor volume, were compared with Wilcoxon rank sum test or Kruskal-Wallis rank sum test (multiple comparisons), using Benjamini-Hochberg procedure for controlling the false discovery rate in multiple comparisons. Spearman rank correlation was used to model the relation between the volume of two ADC-rCBV ROIs and the volume of CE and FLAIR ROIs. We analyzed the available survival data of 80 patients, who received maximal surgery followed by standard regimen of radiotherapy with concomitant temozolomide (TMZ) chemotherapy followed by adjuvant TMZ chemotherapy. Kaplan-Meier and Cox proportional hazards regression analyses were performed to evaluate patient survival. For the Kaplan-Meier analysis, the volumes of ROIs and MRS variables were dichotomized for OS and PFS before the log-rank test using optimal cutoff values calculated by the function of ‘surv_cutpoint’ in R Package “survminer”. Patients who were alive at the last known follow-up were censored. Multivariate Cox regression with forward and backward stepwise procedures was performed for the selection of prognostic variables, accounting for relevant covariates, including IDH-1 mutation, MGMT methylation, sex, age, extent of resection and contrast-enhancing tumor volume. The forward procedure initiated from the model which only contained one covariate. The backward procedure initiated from the model in which all covariates were included. For each step, the model obtained after adding or deleting one covariate was evaluated using the Akaike Information Criterion (AIC). The final multivariate Cox regression model was constructed using only the features selected by the stepwise procedures. The hypothesis of no effect was rejected at a two-sided level of 0.05.

## Results

### Patients

After surgery, 19 (14.5%) patients were excluded due to non-glioblastoma pathological diagnosis and 112 patients (mean age 59.4 years, range 22-76, 84 males; Overview: Table 1) were included. Among them, 82 (73.2%) patients received standard dose of radiotherapy plus temozolomide concomitant and adjuvant chemotherapy post-operatively. Complete resection of contrast enhanced tumor was achieved in 75 of 112 (67.0%) patients. Seven of 112 (6.3%) patients had the IDH-1 R132H mutation and 105 of 112 (93.8%) patients were IDH-1 wild type. MGMT-methylation status was available for 108 patients, among which 48 (44.4%) patients were methylated. Eighty of 112 (71.4%) patients had survival data available. The median progression-free survival (PFS) of these patients was 265 days (range 25-1130 days) and overall survival was 455 days (range 52-1376 days).

**Table 1.**
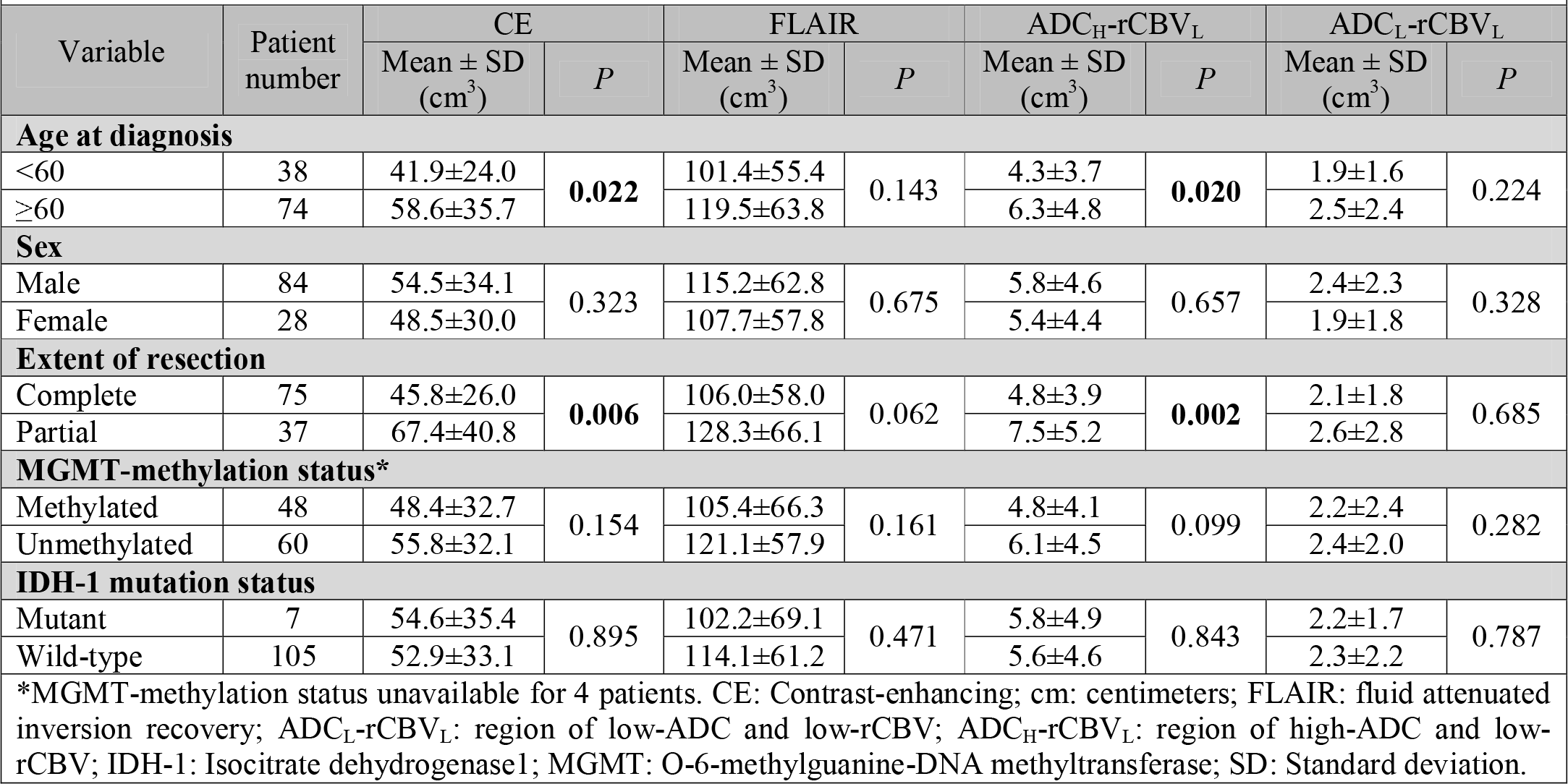
Patient clinical characteristics and ROI volumes

### Multiparametric MRI identifies two low perfusion compartments

The volumes of ROIs for different patient subgroups is shown and compared in Table 1. The interrater variability of the ROIs showed excellent agreement between the two raters. Dice scores of CE regions and FLAIR regions are 0.85 ± 0.10 and 0.86 ± 0.10 respectively. The ADC_H_-rCBV_L_ compartment (volume 5.7 ± 4.6 cm^3^) was generally larger than the ADC_L_-rCBV_L_ compartment (volume 2.3 ± 2.2 cm^3^) (*P* < 0.001). Completely resected tumors had smaller CE volume (*P* = 0.006) and smaller ADC_H_-rCBV_L_ compartment (*P* = 0.002). Figure 4 (A, D) shows the two compartments for two different tumors.

**Figure 4.**
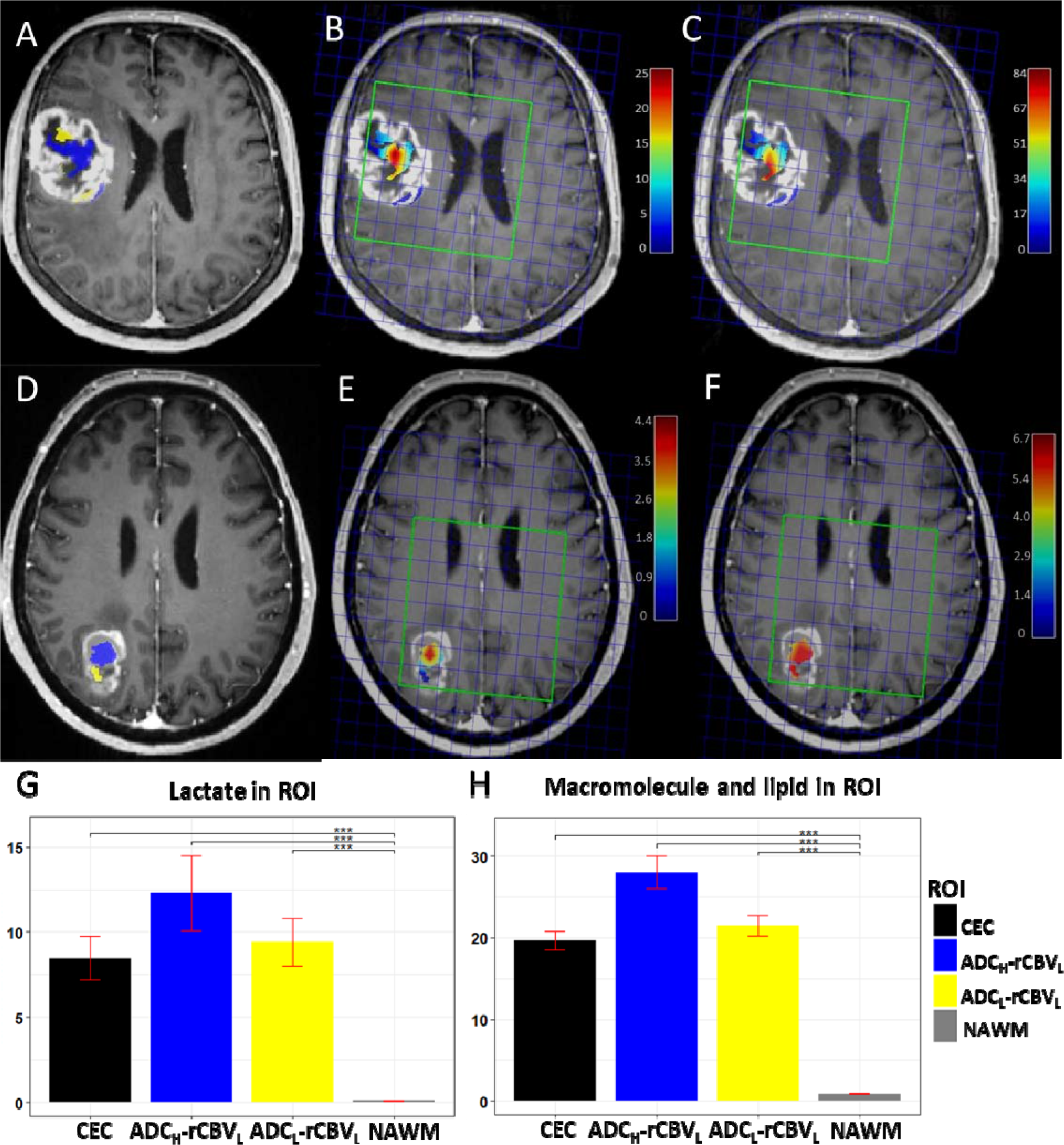
Two hypoxic compartments and MRS characteristics. Case 1: A-C; Case 2: D-F. A & D show the location of ADC_L_-rCBV_L_ (yellow) and ADC_H_-rCBV_L_ (blue) compartments. B & E demonstrate the Lac/Cr ratios of the two compartments. C & F demonstrate the ML9/Cr ratios in the two compartments. The color bar show the level of metabolites (red: high, blue: low). Note that case 1 shows greater tumor volume and higher lactate level. G & H demonstrate the MRS characteristics of the compartments over the patient cohort. Yellow: ADC_L_-rCBV_L_; blue: ADC_H_-rCBV_L_; black: contrast-enhancing control (CEC); grey: normal-appearing white matter (NAWM). G: mean Lac/Cr level; H: mean ML9/Cr. ***: *P* < 0.001.

### Low perfusion compartments displayed hypoxic and pro-inflammatory metabolic signatures

The low perfusion and high diffusion compartment ADC_H_-rCBV_L_ showed a significantly higher lactate/creatine (Lac/Cr) ratio than NAWM (*P* < 0.001), as well as an increased ML9/Cr ratio compared to NAWM (*P* < 0.001). Similarly, the ADC_L_-rCBV_L_ compartment displayed higher Lac/Cr ratio and ML9/Cr ratio than NAWM (both *P* < 0.001). Although not significant, the Lac/Cr and ML9/Cr ratios in the ADC_H_-rCBV_L_ compartment were higher than the ADC_L_-rCBV_L_ compartment (Table S1). Figure 4 shows overlays between the rCBV-ADC compartments and the MRS voxels for two example cases, as well as the metabolite levels in each ROI and compartment.

### Low perfusion compartments exhibited diverse effects on tumor invasion

The contrast-enhancing (CE) tumor volume was significantly correlated with the Lac/Cr ratio in the ADC_L_-rCBV_L_ (*P* = 0.018, rho = 0.34), indicating that this compartment had higher lactate level in tumors with a larger CE volume. Interestingly, the volume of tumor infiltration beyond contrast enhancement, which was delineated on FLAIR images and normalized by CE volume, showed a moderate positive correlation with the proportional volume of the ADC_L_-rCBV_L_ compartment (*P* < 0.001, rho = 0.42) and a negative correlation with the proportional volume of the ADC_H_-rCBV_L_ compartment (*P* < 0.001, rho = -0.32). This indicated that tumors with a large infiltrative volume tended to have smaller ADC_H_-rCBV_L_ and larger ADC_L_-rCBV_L_ compartments. The correlations of ROI volumes are demonstrated in Figure S2.

### The ADC_L_-rCBV_L_ compartment of minimally invasive tumors is less hypoxic

The minimally invasive phenotype displayed a lower volume of ADC_L_-rCBV_L_ compartment than the localized (*P* = 0.031) and diffuse phenotype (not significant), and a higher volume of ADC_H_-rCBV_L_ compartment than the localized (*P* = 0.024) and diffuse phenotype (not significant), suggesting the effects of the two low perfusion compartments to tumor invasiveness were different. Of note, the minimally invasive phenotype displayed lower Lac/Cr ratio compared to the localized (*P* = 0.027) and diffuse phenotype (*P* = 0.044), indicating that the ADC_L_-rCBV_L_ compartment experienced less hypoxic stress in the minimally invasive tumors. A full comparison between the three invasive phenotypes can be found in Table S2.

### Low perfusion compartments exhibited diversity in treatment response

First, we used multivariate Cox regression to analyse all the relevant clinical factors. The results showed that extent of resection (EOR) (PFS: hazard ratio [HR] = 2.825, *P* = 0.003; OS: HR = 2.063, *P* = 0.024), CE tumor volume (OS: HR = 2.311, *P* < 0.001) and FLAIR tumor volume (OS: HR = 0.653, *P* = 0.031) were significantly associated with survivals.

Next, we included the volumes of two compartments and their weighed average Lac/Cr ratios into the survival models. We used a stepwise procedure to select all prognostic variables. The results showed that higher volumes of the two compartments were associated with better PFS (ADC_H_-rCBV_L_: HR = 0.102, *P* = 0.049; ADC_L_-rCBV_L_: HR = 0.184, *P* = 0.033), whilst the higher Lac/Cr ratio in the two compartments was associated with worse PFS (ADC_H_-rCBV_L_: HR = 6.562, *P* = 0.002; ADC_L_-rCBV_L_: HR = 2.995, *P* = 0.047). Further, the higher Lac/Cr ratio in the ADC_L_-rCBV_L_ compartment was also significantly associated with worse OS (HR = 4.974, *P* = 0.005). In contrast, the Lac/Cr ratio in the contrast-enhancing control regions was associated with better survivals (PFS: HR = 0.053, *P* = 0.001; OS: HR = 0.090, *P* = 0.007). The results of the Cox proportional hazards models are described in Table 2 and the Kaplan-Meier survival curves using Log-rank test are shown in Figure 5.

**Figure 5.**
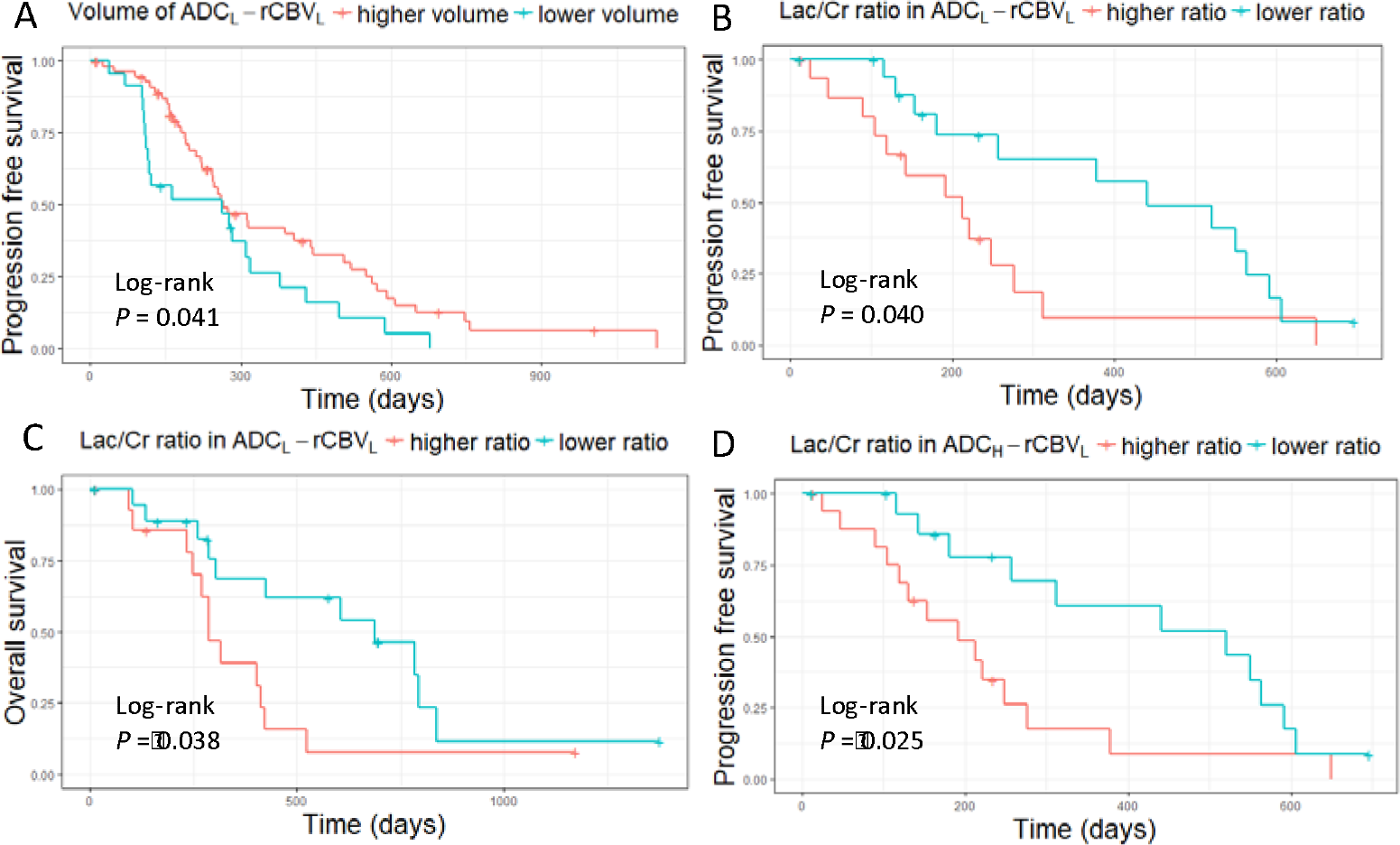
Kaplan-Meier plots of survival analysis. Log-rank tests show larger proportional volume of ADC_L_-rCBV_L_ compartment is associated with better PFS (*P* = 0.041) (A), while higher Lac/Cr ratio in this compartment is associated with worse PFS (*P* =0.040) (B) and OS (*P* = 0.038) (C). Higher Lac/Cr ratio in the ADC_H_-rCBV_L_ compartment is associated with worse PFS (*P* = 0.025) (D).

**Table 2.** Univariate and multivariate modeling of survivals

| Table 2. Univariate and multivariate modeling of survivals |  |  |  |  |  |  |  |  |  |  |  |  |
| --- | --- | --- | --- | --- | --- | --- | --- | --- | --- | --- | --- | --- |
| Factor | PFS |  |  |  |  |  | OS |  |  |  |  |  |
|  | Multivariate |  |  | Stepwise |  |  | Multivariate |  |  | Stepwise |  |  |
|  | HR | 95%CI | P | HR | 95%CI | P | HR | 95%CI | P | HR | 95%CI | P |
| Age | 1.007 | 0.979-1.035 | 0.645 |  |  |  | 1.002 | 0.971-1.033 | 0.911 | 0.927 | 0.860-0.998 | **0.045** |
| Sex (M) | 1.499 | 0.838-2.681 | 0.172 | 5.043 | 1.063-23.91 | **0.042** | 1.252 | 0.662-2.365 | 0.490 |  |  |  |
| extent of resection (partial) | 2.825 | 1.417-5.635 | **0.003** | 4.531 | 1.002-20.49 | **0.050** | 2.063 | 1.099-3.874 | **0.024** | 12.18 | 2.701-54.91 | **0.001** |
| MGMT methylation status* | 0.624 | 0.366-1.063 | 0.083 | 0.392 | 0.125-1.233 | 0.109 | 0.647 | 0.358-1.167 | 0.148 | 0.231 | 0.070-0.762 | **0.016** |
| IDH mutation status | 0.902 | 0.278-2.926 | 0.864 |  |  |  | 0.900 | 0.256-3.170 | 0.870 |  |  |  |
| CE volume | 1.291 | 0.861-1.935 | 0.216 | 6.760 | 0.696-65.63 | 0.099 | 2.311 | 1.527-3.499 | **<0.001** | 3.080 | 1.487-6.383 | **0.002** |
| Flair volume | 0.775 | 0.519-1.157 | 0.212 | 2.008 | 0.818-4.926 | 0.128 | 0.653 | 0.444-0.961 | **0.031** |  |  |  |
| ADC<sub>L</sub>-rCBV<sub>L</sub> volume |  |  |  | 0.184 | 0.039-0.874 | **0.033** |  |  |  |  |  |  |
| ADC<sub>H</sub>-rCBV<sub>L</sub> volume |  |  |  | 0.102 | 0.011-0.992 | **0.049** |  |  |  |  |  |  |
| Lac/Cr in ADC<sub>H</sub>-rCBV<sub>L</sub> |  |  |  | 6.562 | 2.023-21.29 | **0.002** |  |  |  | 2.367 | 0.825-6.790 | 0.109 |
| Lac/Cr in ADC<sub>L</sub>-rCBV<sub>L</sub> |  |  |  | 2.995 | 1.012-8.861 | **0.047** |  |  |  | 4.974 | 1.608-15.39 | **0.005** |
| Lac/Cr in CEC |  |  |  | 0.053 | 0.010-0.295 | **0.001** |  |  |  | 0.090 | 0.016-0.520 | **0.007** |
| *MGMT-methylation status unavailable for 4 patients. PFS: progression-free survival; OS: overall survival; HR: hazard ratio; CI: confidence interval; IDH-1: Isocitrate dehydrogenase 1; MGMT: O-6-methylguanine-DNA methyltransferase; CE: Contrast-enhancing; FLAIR: fluid attenuated inversion recovery; Lac: lactate; ADC<sub>L</sub>-rCBV<sub>L</sub>: region of low-ADC and low-rCBV; ADC<sub>H</sub>-rCBV<sub>L</sub>: region of high-ADC and low-rCBV; CEC: contrast enhanced control. |  |  |  |  |  |  |  |  |  |  |  |  |

## Discussion

This study combined perfusion and diffusion parameters to quantify two low perfusion compartments that may be responsible for treatment resistance. Our findings are consistent with recent studies suggesting various adaptive phenotypes exist under selective pressure^14^ and can predict patient survival.^29^ The proposed approach can visualize and quantify these sub-regions non-invasively using physiological imaging, which may potentially improve on the commonly-used weighted structural imaging.

The clinical values of the individual image derived parameters have been extensively assessed previously and are supported by our findings. Among them, rCBV is associated with tumor angiogenesis and proliferation, and has recently been found to indicate IDH mutation status, associated with hypoxia-initiated angiogenesis.^30^ Decreased ADC is thought to represent higher tumor cellularity and cell packing^31^ and has been associated with shorter survival^32^. However, it remained unknown if integrating these markers could visualize clinically relevant intratumor habitats. We identified two low perfusion compartments with confirmed hypoxic stress demonstrated by elevated lactate levels compared to normal and abnormal controls. With similar perfusion levels in these two compartments, the higher cellularity (ADC_L_) in the ADC_L_-rCBV_L_ compartment suggests that this compartment may display a higher degree of mismatch between supply and demand of nutrients.

We measured the lactate (Lac) and macromolecule and lipid levels at 0.9 ppm (ML9) in the spectra, as an increase in Lac is frequently detected in hypoxic microenvironments, while ML9 was recently associated with pro-inflammatory microglial response to immune stimulus.^33^ The elevated ML9/Cr ratios may suggest both compartments displayed elevated inflammation response,^33^ potentially due to recruitment of tumor-promoting inflammatory cells by necrotic cells.^34,35^ The positive correlation between contrast-enhancing tumor volume and Lac/Cr ratios in the ADC_L_-rCBV_L_ compartment could indicate an acquired hypoxic microenvironment as the solid tumor grows. When evaluating the non-enhancing regions surrounding the solid tumor, we found that tumors with large infiltrative volume tended to have smaller ADC_H_-rCBV_L_ and larger ADC_L_-rCBV_L_ compartments, suggesting that the latter compartment might be more responsible for tumor infiltration. This was supported by our findings that minimally invasive phenotypes displayed significantly lower lactate levels in the ADC_L_-rCBV_L_ compartment.

We further investigated the effects of the two compartments to patient survivals in a cohort who have received standard dose of concurrent temozolomide (TMZ) chemoradiotherapy followed by adjuvant TMZ after surgery. Interestingly, a higher Lac/Cr ratio in the two compartments was related to worse outcomes (HR > 1) while this ratio in other contrast-enhancing tumor regions showed a reduced hazard (HR < 1). This suggests that the resistant phenotypes induced by hypoxia mainly reside in the two identified compartments. As the ADC_L_-rCBV_L_ compartment was associated with tumor infiltration area, diffusion invasiveness, and significantly affected both PFS and OS, this compartment may be more responsible for treatment resistance. Additionally, we found that the higher volume of both compartments was significantly associated with better survivals in the multivariate model, while higher Lac/Cr ratios were associated with worse survivals. These results suggested that the extent of low perfusion, indicated by the volume, and the intensity of hypoxia, indicated by the lactate level, have different clinical implications. Specifically, the higher proportion of the low perfusion compartments may represent a relatively lower proliferative phenotype, while the more intensive hypoxia in these compartment may represent a more aggressive phenotype.

Our findings have clinical significance, as they can help targeting treatment to the selective pressure within tumors. As radiotherapy and chemotherapy can cause selective stress, our identification of possibly resistant target regions could inform the choice of treatment. Additionally, several recent studies postulated that antiangiogenic agents failed to demonstrate consistent response because they can induce the adaptive clones and thus cause treatment resistance.^16,36^ Our findings of the two low perfusion compartments may provide indications for selection of antiangiogenic therapy. Particularly, we suggest more attention might be needed for patients with larger volumes of ADC_L_-rCBV_L_ compartment when considering antiangiogenic agents. The possibility of aggravating hypoxia in this compartment may lead to a more aggressive phenotype.

There are some limitations in our study. While the number of included patients is fairly large, this was a single cohort study. Due to the limited MRS spatial resolution, the multivariate analysis was based on only a subset of the patient cohort. Similarly, the survival analyses were based on the subset of patients who received standard chemoradiotherapy following surgery. The cut-off values defining the two compartments were based on the upper and lower quartiles of the rCBV and ADC distributions, rather than optimizing for threshold specifically. Lastly, although our imaging markers are well validated histologically from other studies,^37^ full biological validation can only be achieved with multi-region sampling of each tumor.

In conclusion, we showed that multiparametric imaging could identify two low perfusion compartments displaying heterogeneity in extent and intensity of hypoxia, and consequently exhibiting diversity in tumor invasion and treatment response. The findings in our study may help optimize the current clinical routine which is mainly based on non-specific conventional imaging. The compartment demonstrating both low perfusion and restricted diffusion may indicate a habitat resistant to adjuvant therapies, which may aggravate the hypoxic stress. This could provide crucial information for treatment choice in a personalized treatment setting. As our analyses were based on clinically available imaging modalities, this approach could easily be implemented, and potentially extended to other system.

---

**Table S1:** Metabolic characteristics measured by MRS.

| Lac/Cr |  |  |  |  |  |
| --- | --- | --- | --- | --- | --- |
| ROI | Descriptive |  | ADC<sub>L</sub>-rCBV<sub>L</sub> | CEC | NAWM |
| | Mean ± SD | 95% CI | *P* | *P* | *P* |
| ADC<sub>H</sub>-rCBV<sub>L</sub> | 12.3±15.4 | 7.9-16.7 | 0.427 | 0.411 | < 0.001 |
| ADC<sub>L</sub>-rCBV<sub>L</sub> | 9.4±9.8 | 6.6-12.2 | / | 0.407 | < 0.001 |
| CEC | 8.5±8.9 | 5.9-11.0 | / | / | < 0.001 |
| NAWM | 0.10±0.28 | 0.02-0.17 | / | / | / |
| ML9/Cr |  |  |  |  |  |
| ROI | Descriptive |  | ADC<sub>L</sub>-rCBV<sub>L</sub> | CEC-ROI | NAWM-ROI |
| | Mean ± SD | 95% CI | *P* | *P* | *P* |
| ADC<sub>H</sub>-rCBV<sub>L</sub> | 28.0±71.6 | 9.2-46.9 | 0.650 | 0.529 | < 0.001 |
| ADC<sub>L</sub>-rCBV<sub>L</sub> | 21.5±31.2 | 13.3-29.7 | / | 0.492 | < 0.001 |
| CEC | 19.7±29.3 | 12.0-27.4 | / | / | < 0.001 |
| NAWM | 0.87±0.68 | 0.70-1.05 | / | / | / |

ROI: region of interest; ADC<sub>L</sub>-rCBV<sub>L</sub>: ROI of low-ADC and low-rCBV; ADC<sub>H</sub>-rCBV<sub>L</sub>: ROI of high-ADC and low-rCBV; CEC: contrast enhancement control; NAWM: normal appearing white matter; Lac: lactate; ML9: Macromolecule and lipid at 0.9ppm; Cr: creatine; CI: confidence interval.

**Table S2:** Comparison of the three DTI based invasive phenotypes.

|  |  | Diffuse | Localised | Minimal | Comparisons |  |  |
| --- | --- | --- | --- | --- | --- | --- | --- |
|  |  | 10 (15.6%) | 40 (62.5%) | 14 (21.9%) | Localised-Diffuse | Minimal-Diffuse | Minimal-Localised |
| ROI | Variable | Mean ± SD | Mean ± SD | Mean ± SD | *P* | *P* | *P* |
| CE | Volume (cm<sup>3</sup>) | 53.8±35.0 | 39.7±16.0 | 52.7±36.2 | 0.388 | 0.406 | 0.390 |
| FLAIR | Volume (cm<sup>3</sup>) | 116.7±57.3 | 82.0±31.5 | 85.2±68.4 | 0.067 | 0.062 | 0.490 |
| ADC<sub>H</sub>-rCBV<sub>L</sub> | Volume<sup>#</sup> | 0.10±0.04 | 0.08±0.03 | 0.12±0.04 | **0.036** | 0.140 | **0.024** |
| ADC<sub>L</sub>-rCBV<sub>L</sub> | Volume<sup>#</sup> | 0.04±0.03 | 0.06±0.03 | 0.03±0.02 | 0.080 | 0.104 | **0.031** |
| ADC<sub>H</sub>-rCBV<sub>L</sub> | Lac/Cr | 12.9±20.3 | 12.9±12.3 | 3.5±3.8 | 0.206 | 0.182 | 0.190 |
| ADC<sub>L</sub>-rCBV<sub>L</sub> | Lac/Cr | 9.0±9.1 | 10.0±8.9 | 1.8±2.3 | 0.407 | **0.044** | **0.027** |
| CEC | Lac/Cr | 8.8±9.2 | 8.8±7.6 | 2.7±2.7 | 0.218 | 0.080 | 0.073 |
| * raw volumes; <sup>#</sup> proportional volumes; ROI: region of interest; CE: contrast enhancement; cm: centimeters; FLAIR: fluid attenuated inversion recovery; ADC<sub>L</sub>-rCBV<sub>L</sub>: ROI of low-ADC and low-rCBV; ADC<sub>H</sub>-rCBV<sub>L</sub>: ROI of high-ADC and low-rCBV; CEC: contrast enhancement control; Lac: lactate; Cr : creatine ; SD: Standard deviation. |  |  |  |  |  |  |  |

**Figure S1.**
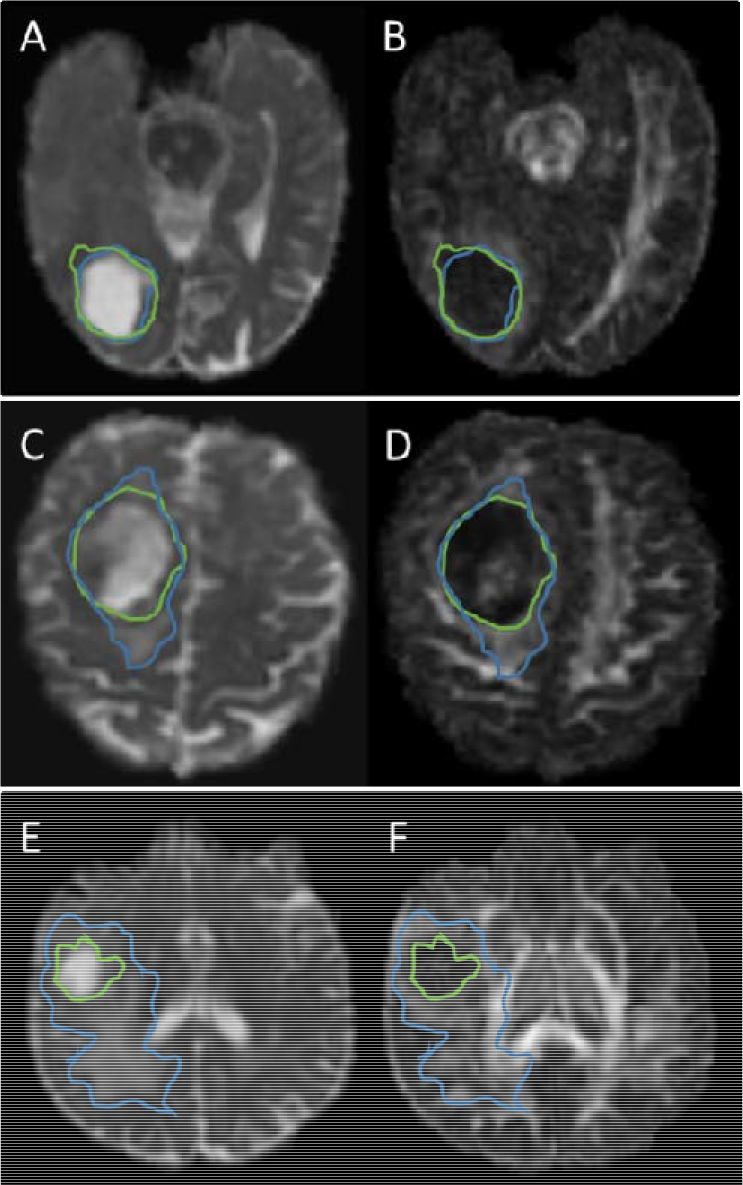
Examples of DTI invasive phenotypes. (A), (C), (E): DTI-p maps with abnormality outlined by the blue line; (B), (D), (F): DTI-q maps with abnormality outlined by the green line. (A) & (B) show a minimal invasive phenotype. The isotropic abnormality is similar to the anisotropic abnormality. (C) & (D) show a localized invasive phenotype. The isotropic abnormality is larger than the anisotropic abnormality in one direction. (E) & (F) show a diffuse invasive phenotype. The isotropic abnormality is larger than the anisotropic abnormality in more than one direction.

**Figure S2.**
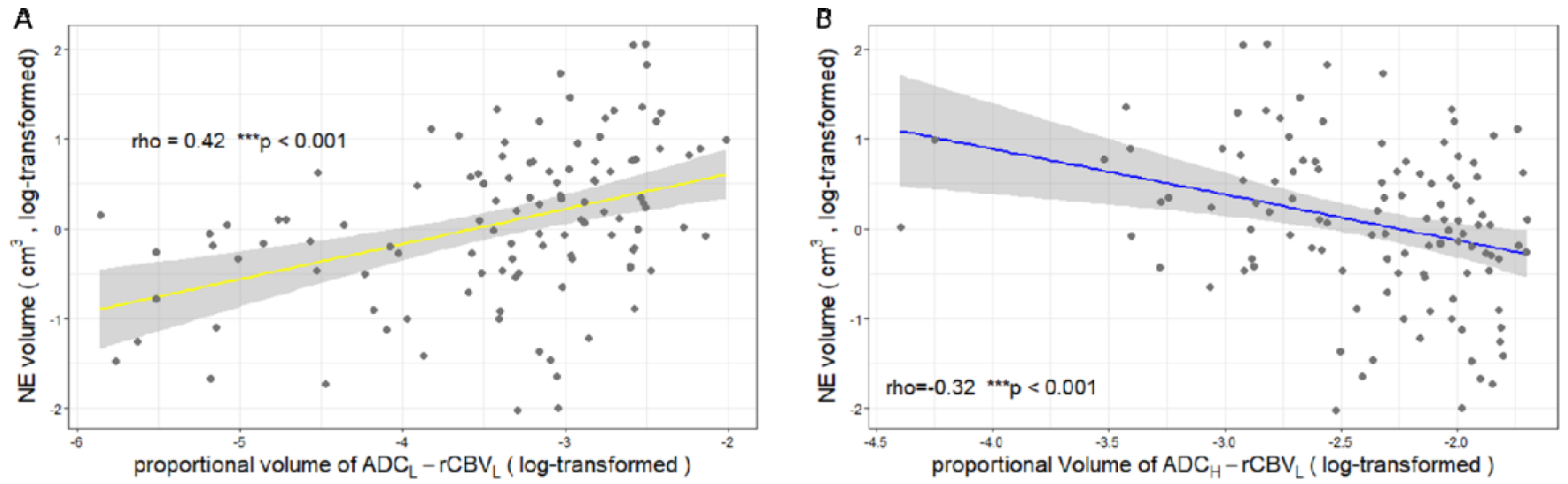
Correlations between two compartments with contrast enhanced tumor and tumor infiltration. Non-enhancing (NE) tumor (measured by the volume of FLAIR beyond contrast enhancement) showed a moderate positive correlation with the proportional volume (log-transformed) of the ADC_L_-rCBV_L_ compartment (A) and negative correlation with the proportional volume (log-transformed) of the ADC_H_-rCBV_L_ compartment (B). ***: p < 0.001.

